# NLRP3 inflammasome is dispensable in methicillin resistant *Staphylococcus aureus* urinary tract infection

**DOI:** 10.1101/2022.11.11.516235

**Authors:** Santosh Paudel, Kenneth A Rogers, Rahul Kumar, Yogesh Saini, Sonika Patial, Ritwij Kulkarni

## Abstract

NLRP3 inflammasome is a cytoplasmic complex that senses molecular patterns from pathogens or damaged cells to trigger an innate immune defense response marked by the production of proinflammatory cytokines IL-1β and IL-18 and an inflammatory death called pyroptosis. The NLRP3 inflammasome is activated in the urinary tract by a variety of infectious and non-infectious insults. In this study, we investigated the role of NLRP3 inflammasome by inducing methicillin resistant *Staphylococcus aureus* (MRSA) ascending UTI in WT and *Nlrp3*^-/-^ mice. At 24 and 72 hpi, compared to the WT, the MRSA-infected *Nlrp3*^-/-^ showed ∼100-fold lower median CFUs, although this reduction was not statistically significant. The ablation of NLRP3 did not affect MRSA-induced urinary immune defenses as indicated by the similar levels of pro-inflammatory cytokines and chemokines and the similar numbers of granulocytes in the bladder and the kidneys of WT and *Nlrp3*^-/-^ mice at 24 h after MRSA infection. However, MRSA-infected *Nlrp3*^-/-^ bladders, but not kidneys, showed significantly higher monocyte infiltration. The histopathological analysis of bladder and kidney sections showed similar inflammation in MRSA-infected *Nlrp3*^*-/-*^ and WT mice. Overall, these results suggest that MRSA-induced urinary NLRP3 activity is dispensable to the host.

**Importance:** Indwelling urinary catheter usage increased susceptibility to methicillin-resistant *Staphylococcus aureus* (MRSA) urinary tract infections (UTI) which can be difficult to treat and can result in potentially fatal complications such as bacteremia, urosepsis, and shock. In this work, we examined the role of NLRP3 inflammasome in MRSA uropathogenesis. In comparison to the WT, mice deficient in NLRP3 activity showed similar MRSA burden and similar inflammation in the bladder and kidney tissues at 24 h after the experimental induction of ascending UTI. These results suggest that NLRP3 inflammasome is not involved in shaping urinary immune defenses during acute MRSA-UTI.

## Introduction

*Staphylococcus aureus* is an atypical cause of asymptomatic bacteriuria and complicated urinary tract infections (UTI) primarily affecting the individuals with indwelling urinary catheters, the elderly, and the hospitalized (1-7). The urinary colonization by *S. aureus* is a major clinical concern because it can lead to life-threatening invasive infections such as bacteremia, urosepsis, and shock (6, 8-10) and because of the increased prevalence of methicillin resistant *Staphylococcus aureus* (MRSA) in urine specimens in the last two decades (6, 10, 11). Previous reports have described specific host-pathogen effectors crucial for MRSA urinary survival and persistence. For example, MRSA infection was reported to augment the catheter implant-mediated localized pro-inflammatory cytokine response and fibrinogen release in the mouse urinary bladder (12). We have reported that *in vitro* exposure to human urine for 2 h increases MRSA virulence and induces metabolic changes necessary for survival in the nutrient-limiting urinary tract (13). However, the role of NLRP3 (NOD (nucleotide oligomerization domain) LRR (leucin-rich repeat) containing receptor, pyrin domain containing protein 3) inflammasome activity in MRSA uropathogenesis has not been determined.

In response to a variety of bacterial molecular patterns, the NLRP3 forms a cytoplasmic inflammasome complex with ASC (apoptosis-associated speck-like protein containing CARD) adaptor, and caspase-1, which cleaves pro-IL-1β and pro-IL-18 into active IL-1β and IL-18 and activates pro-inflammatory programmed cell death called pyroptosis via cleavage of gasdermin D (14). Uropathogenic *Escherichia coli* (UPEC) α-hemolysin has been reported to activate NLRP3-IL1β signaling axis and pyroptosis in macrophages, neutrophils, renal fibroblasts, and bladder epithelial cells (15-21). Being partly responsible for the exfoliation of bladder epithelium and subsequent elimination of adherent and intracellular UPEC, the NLRP3-pyroptosis is an effective immune defense against acute cystitis (15, 22). In contrast to this protective role during acute cystitis, however the NLRP3 activity can also promote chronic UPEC-UTI as the exfoliation exposes the underlying epithelium for UPEC (23). In the case of MRSA, the toxins Panton-Valentine leukocidin (PVL), leukocidin AB (LukAB), and α-hemolysin (Hla) have been reported to trigger NLRP3 inflammasome in myelocytes (24-27). The impact of NLRP3 activity on the outcomes of *S. aureus* infection, however are reported to be dependent on the site of infection. For example, *S. aureus*-induced NLRP3-IL1β signaling axis has been found to be protective during skin and soft tissue infection (28), and detrimental for the host during severe *S. aureus* pneumonia (24). In contrast, NLPR3 protein is not required for the survival in acute central nervous system *S. aureus* infection because in this model AIM2 (absent in melanoma 2) has been reported to replace NLRP3 in the IL-1β processing inflammasome complex (29). The role of NLRP3 in MRSA uropathogenesis has not been deciphered.

We hypothesized a protective role for the NLRP3 inflammasome activity in MRSA acute cystitis. To address this hypothesis, we experimentally induced ascending UTI by inoculating MRSA into the urinary bladders of 8-10 weeks old, female WT and *Nlrp3*^*-/-*^ mice. We observed lower MRSA CFUs in the bladder and the kidneys of *Nlrp3*^*-/-*^ mice at 24 and 72 hpi compared to the WT, although this reduction was not statistically significant. The WT and *Nlrp3*^-/-^ mouse cohorts with MRSA ascending UTI also showed similar levels of pro-inflammatory cytokines (measured by ELISA) and granulocyte infiltration (flow cytometry and histopathology) in the bladder and kidney tissues, although MRSA-infected *Nlrp3*^-/-^ bladders showed significantly higher monocyte recruitment compared to their WT counterparts. The treatment of WT mice with NLRP3 inhibitor, MCC950 at 4 h after MRSA infection resulted in a modest but statistically significant reduction in the bladder burden at 24 hpi compared to the vehicle-treated controls. In summary, the activation of NLRP3 inflammasome appears to be dispensable during MRSA acute UTI with the caveat that our experiments do not examine the effects of MRSA-induced urinary NLRP3 activity on the exfoliation of uroepithelium or on the outcomes of MRSA chronic UTI.

## Materials and Methods

### Bacteria, mice, and the reagents

Uropathogenic MRSA strain, MRSA1369 was grown overnight at 37°C and 200 rpm shaking in tryptic soy broth (TSB). The bacteria from the overnight culture were diluted 1:10 in fresh TSB, incubated at 37°C, 200 rpm shaking to a mid-log phase (OD_600_ = 0.6). To prepare inoculum for the mouse infection, mid-log phase MRSA 1369 culture was washed once in sterile D-PBS (Dulbecco’s phosphate buffered saline) and adjusted to 10^9^ CFU (colony forming unit)/ml. *Nlrp3*^-/-^ mice (Stock #021302) were purchased from The Jackson Laboratory (30). NLRP3 inhibitor MCC950 was purchased from InvivoGen and resuspended in DMSO vehicle before administration at the desired concentration. Other reagents were purchased from Fisher Scientific.

### Mouse model of ascending UTI with and without a catheter implant

C57BL6 WT and *Nlrp3*^*-/-*^ breeding trios from the Jackson Laboratory were housed at the biology department mouse facility located at University of Louisiana at Lafayette. To normalize individual gut microbiota of WT and *Nlrp3*^*-/-*^ mice and to minimize cage differences, we used bedding transfer where the soiled bedding from the cages containing WT and *Nlrp3*^*-/-*^ mice was mixed and distributed equally over a period of three weeks, from weaning till the experimental induction of ascending UTI (31). As approved by the Institutional Animal Care and Use Committee (IACUC) at UL Lafayette (2018-8717-011), we administered via transurethral catheterization 50 μl MRSA 1369 (equivalent to 5 × 10^7^ CFU) into the urinary bladders of anesthetized, 8-10 weeks-old female WT and *Nlrp3*^*-/-*^ mice (32). For the mouse model of catheter-associated UTI (CAUTI), MRSA 1369 inoculum was administered immediately after implanting a 4-to 5-mm piece of silicone catheter (12). Mice were sacrificed at 6, 24, and 72 hours post infection (hpi). The bladder, kidney, and spleen were dilution plated on CHROMagar™ or TS agar to determine organ-specific MRSA burden. Tissue samples were also processed for ELISA, flow cytometry, or histopathology as described elsewhere.

### MCC950 treatment

In separate experiments, at 4 h after induction of ascending UTI, one group of MRSA 1369-infected WT littermates were intraperitoneally injected with 10mg/kg MCC950 while the control group was injected with DMSO vehicle. Mice were sacrificed 24 hpi and MRSA burden in the bladder, the kidneys, and the spleen was determined by dilution plating tissue homogenates on CHROMagar™ or TS agar plates.

### Cytokine profiling by ELISA

The bladder and kidney homogenates in sterile D-PBS were filtered through 0.65 μm Ultrafree®-MC Centrifugal Filter (Millipore sigma) and the total protein concentration was estimated using Pierce BCA protein assay kit (Thermo-scientific). The levels of cytokines IL-1β, IL-6, IL-10, IL-17A, TNF-α, CXCL1 (KC), CCL2 (MCP-1), CCL3 (MIP1α), CCL5 (RANTES), and IFN-γ, in the tissues homogenates were estimated using MILLIPLEX^®^ Mouse Cytokine/Chemokine Magnetic Bead Panel (MCYTOMAG-70K-10C). Each cytokine in an individual mouse tissue was presented as a scattered diagram showing the amount of cytokine/g of total protein with median as the central tendency.

### Immune cell infiltration in the bladder and the kidney tissues

Specific immune cells infiltrating the bladder and the kidneys of WT and *Nlrp3*^-/-^ mice were identified using a panel of fluorescent-labelled antibodies for flow cytometry (Table 2 and (33)). Prior to the antibody treatment, the chopped organs were enzymatically digested in RPMI medium containing collagenase IV (8 mg/ml for the bladder and 2 mg/ml for the kidney) and DNase I (1 μl) at RT for 90 min, 250 rpm shaking with frequent pipetting to mix. The cell suspension was passed through a 35 μm filter strainer (Falcon®) to remove leftover tissue pieces, and washed once in D-PBS(2000 rpm, 5 min, RT).

After treatment with RBC lysis buffer (RT, 10 min), the cells were centrifuged. Next, the cell pellets were stained (in tubes protected from light) with 1 μl live/dead marker (Alexa Fluor 430 NHS Ester (Succinimidyl Ester), ThermoFisher) RT, 25 min, 2 μl of Fc block (surface staining, 4°C, 10 min), and then with an antibody cocktail (2μl/antibody, RT, 15 min). Between the two staining steps, the cell pellets were washed once in FACS buffer (D-PBS + 2%FBS). After the final staining step, the cells were resuspended in 250 μl fixation buffer (4°C, 20 min), washed once in FACS buffer, and resuspended in FACS buffer for use in flow-cytometry. The data were analyzed with FlowJO™ version 10. After gating on CD45^+^ cells, we detected monocytes (MHCII^-^CD11b^+^ Ly6G^-^), neutrophils (MHCII^-^CD11b^+^ Ly6G^+^), eosinophils (MHCII^-^CD11b^+^ SiglecF^+^Ly6G^-^), and mast cells (CD117^+^) (Fig S3).

### Histopathological examination of bladder and kidney

WT and *Nlrp*3^-/-^ mouse bladders and kidneys (from MRSA infected and control mice) were preserved in 10% formalin, embedded in paraffin, sectioned, and stained with hematoxylin and eosin. A veterinary pathologist assessed sections of bladder and kidney microscopically in a blinded manner following previously determined semiquantitative scoring scheme to score the severity and extent of inflammation (34). For bladder sections, widespread inflammation, thrombosed vessels, and marked submucosal edema were assigned a score of 3; mixed inflammation in mucosa and submucosa and around vessels, a score of 2; scattered neutrophils in submucosa and migrating through mucosa, a score of 1; while normal sections were assigned a score of 0. For kidney sections, many neutrophils in pelvic lumen and within the tissue were assigned a score of 3; clustered neutrophils in pelvic lumen; and inflammation within the epithelium and surrounding stroma, a score of 2; scattered neutrophils migrating though pelvic epithelium, a score of 1; while normal sections were assigned a score of 0.

### Statistical analysis

Statistical tests were performed using Prism 9.4.1 (www.graphpad.com). Data from multiple biological replicates with two or more technical replicates for each experiment were pooled together. Error bars in the figures represent standard deviation. Organ burden, cytokine amount, number of infiltrating immune cells, and histological scores between the WT and the *Nlrp*3^-/-^ mice were compared using Mann-Whitney U statistic. Data were considered statistically significant if P ≤ 0.05.

## Results

### The kinetics of MRSA colonization in WT and *Nlrp3*^*-/-*^ mice

The ability of uropathogenic MRSA 1369 to infect the urinary tract and to disseminate to the spleen was compared in C57BL6 WT and *Nlrp3*^*-/-*^ mouse models of ascending UTI by enumerating CFU burden in bladder, kidneys, and spleen at 6 (Fig 1A), 24 (Fig 1B), and 72 (Fig 1C) hpi. At 6 hpi, we observed similar bacterial burden in the bladder and kidney tissues from WT and *Nlrp3*^-/-^ mice (Fig 1A). At 24 hpi, however, the *Nlrp*3^-/-^ mice showed ∼17-fold reduction in the median bladder CFUs (P= 0.22) and 8-fold reduction in the median kidney CFUs (P= 0.12) compared to the WT. At 72 hpi, compared to the WT the *Nlrp3*^-/-^ mice showed ∼6-fold reduction in median bladder CFU (P= 0.3) and ∼9-fold reduction in median kidney CFUs (P= 0.31). Thus, compared to the WT, the *Nlrp3*^-/-^ mice showed consistent but statistically insignificant reduction in the kidney and bladder CFU burdens at 24 or 72 hpi. We also observed modest reduction in the number of *Nlrp3*^-/-^ mice showing MRSA dissemination to spleen compared to their WT counterparts. For example, we detected MRSA in the spleen homogenates of 1/6 *Nlrp3*^-/-^ and 3/6 (50%) WT mice at 6 hpi, 0/18 *Nlrp3*^-/-^ and 3/14 (21%) WT mice at 24 hpi. MRSA CFUs were not detected in the spleen homogenates of either WT or *Nlrp3*^-/-^ mice at 72 hpi.

**Figure 1:**
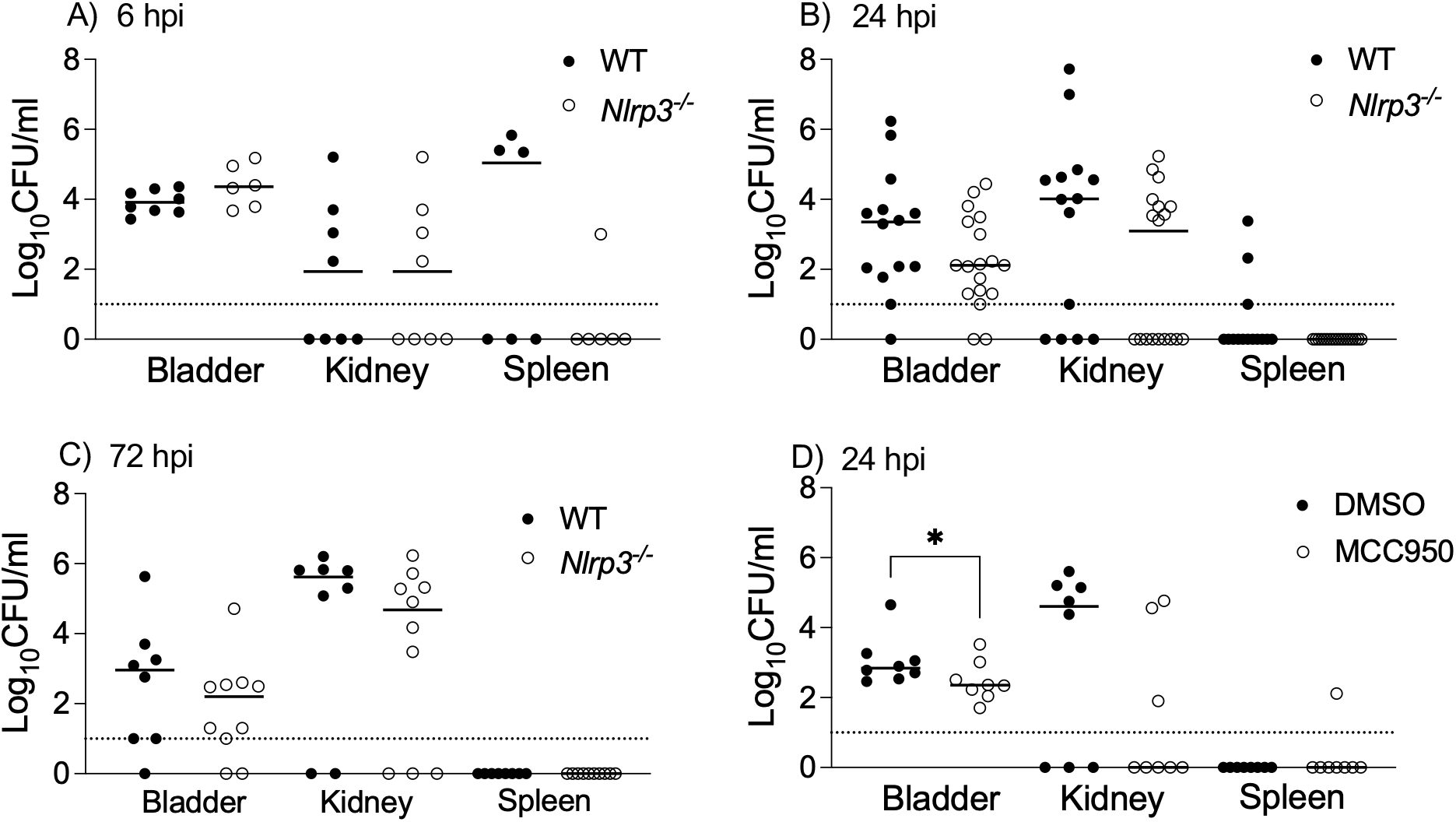
MRSA UTI in WT and Nlrp3^-/-^ mice. Female WT (control) and *Nlrp3*^−/−^ mice were inoculated transurethrally with 5 × 10^7^ CFU of uropathogenic MRSA strain, MRSA 1369. Mice were sacrificed and the bacterial burden in the bladder, the kidneys, and the spleen was determined at 6 hpi (A), 24 hpi (B), and 72 hpi (C). In separate experiments, WT C57BL6 mice infected with MRSA1369 were injected 4hpi intraperitoneally with either 10mg/kg MCC950 or DMSO vehicle. MRSA CFUs in the bladder, the kidneys and the spleen were determined at 24hpi (D). Scatter plots show CFU counts from a single mouse (n= 6 to 18/ group) with the median as the central tendency; the dotted lines show the limit of detection. Statistical significance was determined by Mann-Whitney U test. For all figures, P≤0.05 was considered significant and indicated by *.

Furthermore, we observed similar median CFUs in the bladder, kidney, or spleen homogenates of female WT and *Nlrp3*^-/-^ mice with MRSA CAUTI at 24 hpi (Fig S1A) as well as 72 hpi (Fig S1B). The median bacterial CFUs recovered from the silicone catheters implanted in the bladder at the time of infection were also similar between WT and *Nlrp3*^-/-^ mice at 24 and 72 hpi (Fig S1).

Next, we treated MRSA-infected WT mice with an NLRP3 inhibitor, MCC950 at 4 h after the induction of ascending UTI without catheter implants. At 24 hpi, we observed that the MCC950-treated mice had ∼3-fold reduced median bladder bacterial burden (P=0.049) compared to DMSO vehicle-treated control mice (Fig 1D). The MCC950 treatment did not affect median kidney bacterial burden significantly (MCC950-treated median <LOD, DMSO-treated median= 40,500 CFU/ml, P=0.2), although it must be noted that we detected kidneys CFUs in fewer MCC950-treated versus DMSO-treated mice (3/8 versus 5/8, P= 0.6 by Fisher’s exact test). The spleen CFUs were detected only in one MCC950-treated mouse (Fig 1D). The reduction in the MRSA burden in the MCC950-treated mouse bladder was not due to bacterial killing by MCC950 as confirmed by similar CFUs observed in MRSA 1369 cultures in the presence of MCC950 or DMSO vehicle control (Supplementary Figure S2) Overall, these results indicate that the genetic ablation of NLRP3 is dispensable for MRSA UTI either with or without the catheter implant and that the pharmacological inhibition of NLRP3 inflammasome activity 4 hours after experimental induction of MRSA UTI modestly reduces bladder MRSA burden.

### MRSA-induces similar levels of cytokine production in the WT and *Nlrp3*^-/-^ mice

Next, we assayed the effects of *Nlrp3* ablation on MRSA-induced production of pro-inflammatory cytokines (IL-1β, IL-6, IL-17A, and TNF-α) and chemokines (CCL2, CCL3, CCL5, CXCL1, and CXCL2) and anti-inflammatory cytokine IL-10 at 24 hpi time point at which MRSA-infected bladder tissue show significant localized inflammation (12). MRSA infection induced production of IL-6, CXCL1, CCL2, and CCL3 in mouse bladders (Fig 2A) and IFNγ, CXCL1, and IL-10 in mouse kidneys (Fig 2B). Although, compared to the WT, none of the cytokines was produced in significantly different amounts by MRSA-infected *Nlrp3*^-/-^ mice either in the bladder (Fig 2A) or the kidney (Fig 2B) tissues.

**Figure 2:**
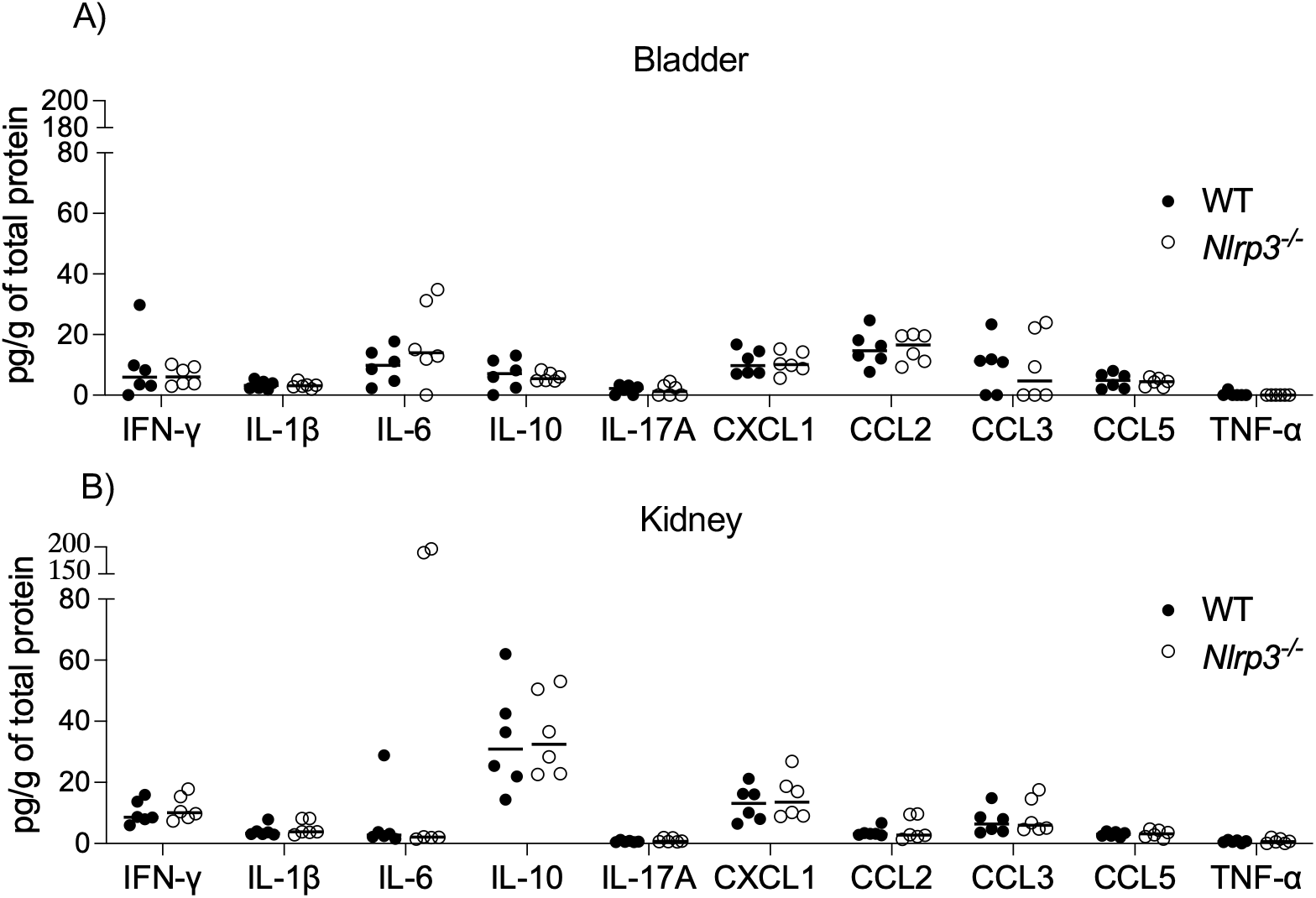
Cytokine profiling of MRSA-infected WT and Nlrp3^-/-^ mouse urinary tracts. MRSA 1369-infected female WT (control) and *Nlrp3*^−/−^ mice were sacrificed at 24 hpi. Cytokines and chemokines (IL-1β, IL-6, IL-10, IL-17A, TNF-α, CXCL1 (KC), CCL2 (MCP-1), CCL3 (MIP1α), IFN-γ, CCL5 (RANTES) produced in bladder (A) and kidneys (B) were quantified by Multiplex-ELISA. The data were compared using Mann-Whitney U test.

### The ablation of *Nlrp3* does not affect MRSA-induced immune cell infiltration in mouse bladder and kidney tissues

Next, we used flow cytometry to characterize the granulocytes and monocytes infiltrating the bladder and the kidneys of MRSA 1369-infected WT and *Nlrp3*^-/-^ mice at 24 hpi. The fluorescent antibodies used to stain specific cell surface markers are presented in Table 1. The gating strategy to differentiate between different immune cells is presented in supplementary figure S3. Among the CD45^+^ cells, the dominant cell populations were neutrophils (MHCII^-^/CD11b^+^/Ly6G^+^), monocytes (MHCII^-^/CD11b^+^/Ly6G^-^), eosinophils (MHCII^-^/CD11b^+^/SiglecF^+^/Ly6G^-^), and mast cells (CD117^+^). While MRSA-infected *Nlrp3*^-/-^ bladders showed higher number of monocytes, eosinophils, and mast cells compared to those in MRSA infected WT bladder tissues (Fig 3A, B), this increase was statistically significant only in the case of monocytes. Compared to their WT counterparts, MRSA-infected *Nlrp3*^-/-^ kidneys showed statistically insignificant reduction in neutrophils population, while the monocytes, eosinophils, and mast cells populations were unaffected (Fig 3C, D).

**Table 1:** List of antibodies used for flowcytometry.

| Antibodies | Conjugate | Clone |
| --- | --- | --- |
| CD64 | BV 786 | X54-5/7.1 |
| SiglecF | BV711 | M290 |
| CD3 | BV650 | 145-2C11 |
| CD45 | BV605 | 30-F11 |
| Live/Dead | Alexa fluor 430 |  |
| Fc epsilon RI | Pac Blue | MAR1 |
| Ly6C | PE-Cy7 | HK1.4 |
| CD11c | PerCp-Cy5.5 | N418 |
| CD103 | PE-CF594 | M290 |
| c-kit (CD117) | PE | 2B8 |
| CD11b | FITC | M1/70 |
| MHC-II (I-A/I-E) | APC-Fire750 | M5/114.15.2 |
| F4/80 | Alexa Fluor 700 | CI:A3-1 |

**Figure 3:**
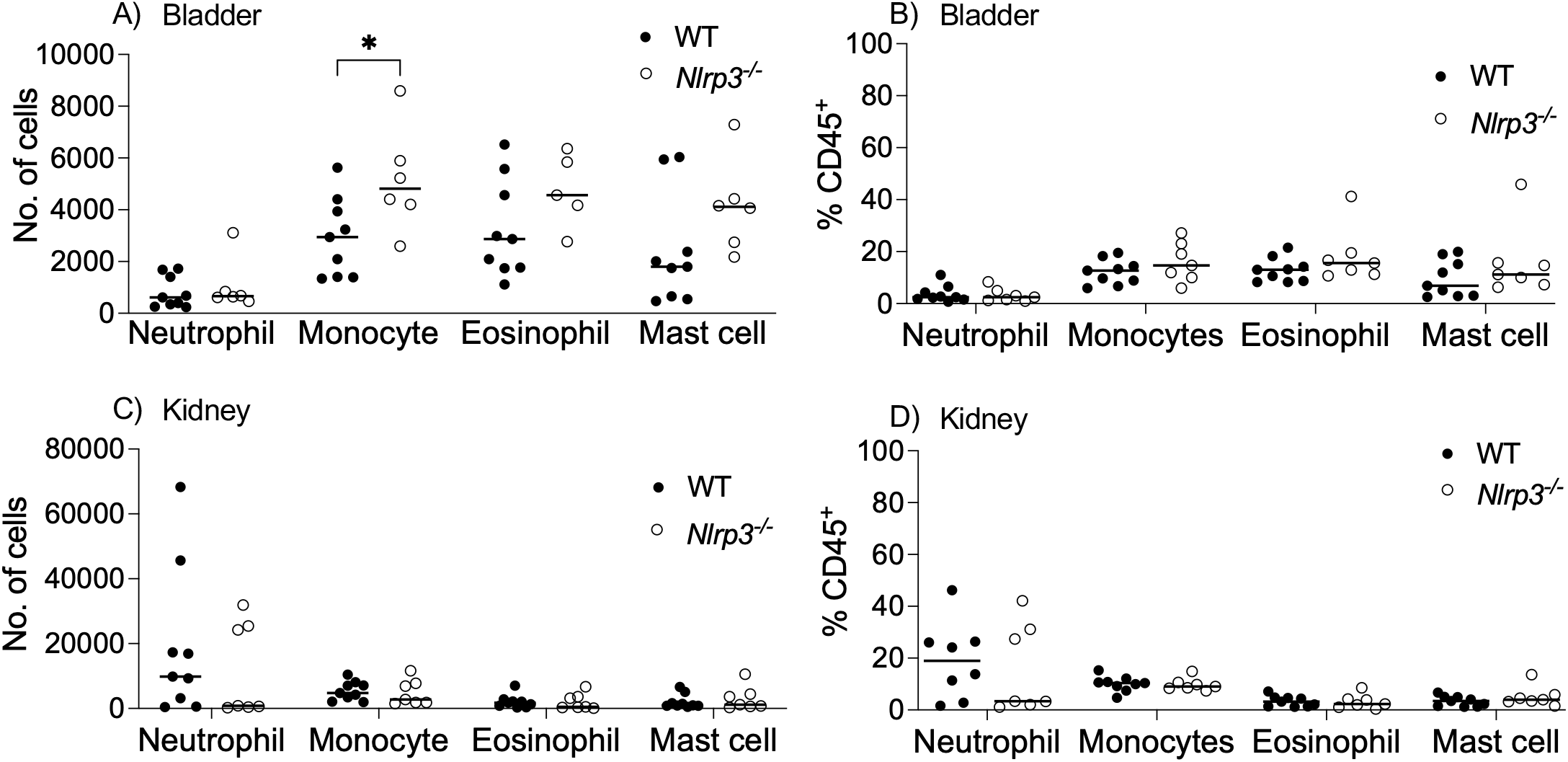
Immune cell infiltration to MRSA-infected WT and Nlrp3^-/-^ mouse urinary tracts. The bladder (A, B) and kidney (C, D) homogenates from WT and *Nlrp3*^*−/−*^ mice infected with MRSA 1369 for 24 h were analyzed by flow cytometry using the gating strategy for CD45^+^ lymphocytes provided in Fig S3. Specific lymphocyte types are shown as the total number of cells (A, C) and the percentage of CD45^+^ lymphocytes (B, D). Statistical significance was determined by Mann-Whitney U test.

### Histopathological examination of WT and *Nlrp3*^-/-^ bladder and kidney sections

In comparison to the PBS inoculated controls, MRSA-infected WT as well as *Nlrp3*^-/-^ mice showed significantly higher presence of inflammatory cells in bladder (Fig 4 A) and kidney samples (Fig 4 C). Next, we scored these tissues for the signs of inflammation on a scale of 1-3 and individual scores for each tissue were plotted. We observed that MRSA infection significantly increased the median inflammation scores in bladder (Fig 4B) and kidney (Fig 4D) of both WT and *Nlrp3*^*-/-*^ mice compared to PBS-inoculated controls. However, within MRSA-infected bladder or kidney tissues, the severity of inflammation between WT and *Nlrp3*^*-/-*^ mice was similar.

**Figure 4:**
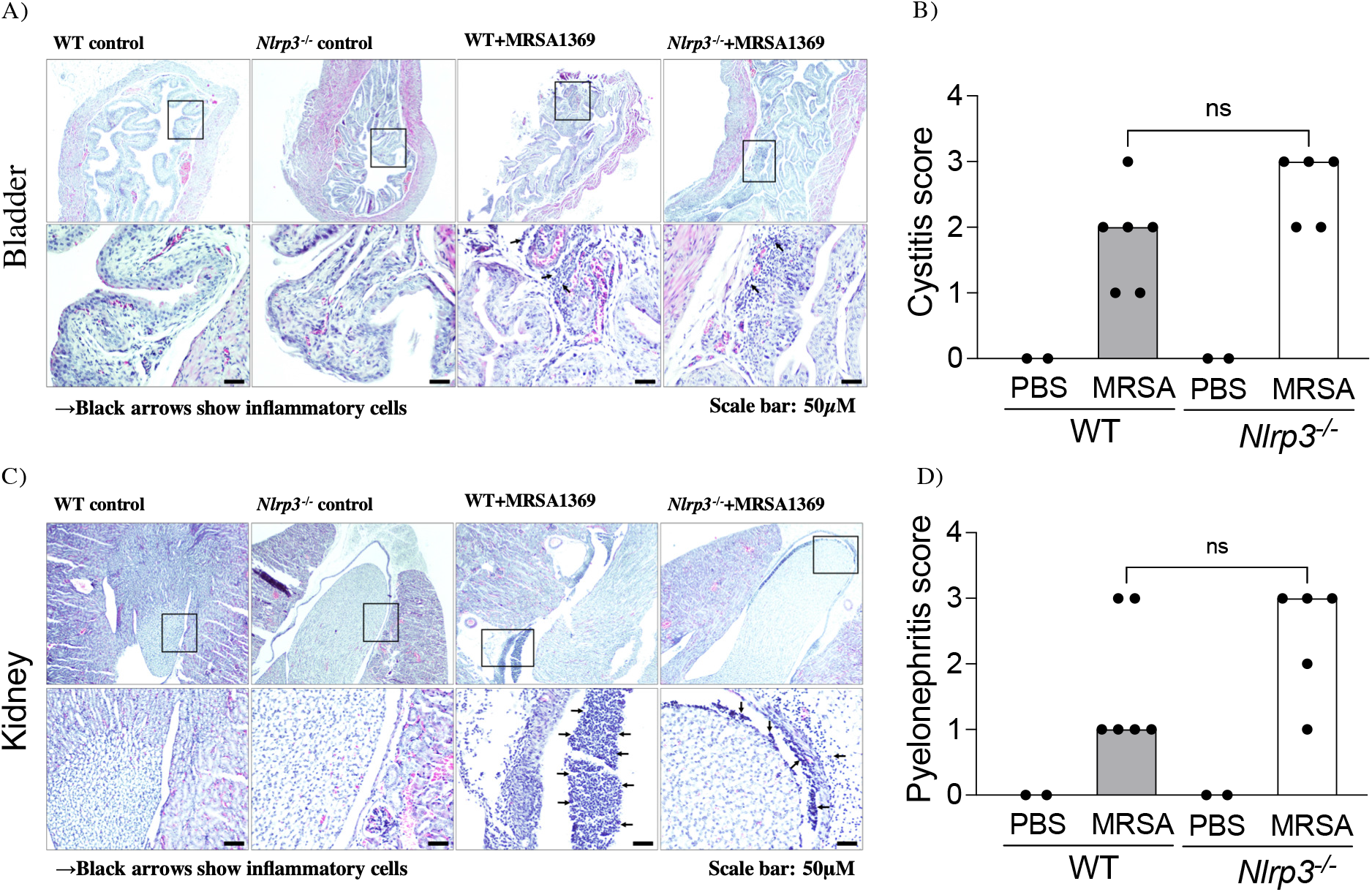
Histopathological examination of MRSA-infected WT and Nlrp3^-/-^ bladder and kidney sections. Bladder (A) and kidney (C) sections from control (PBS) and MRSA-infected WT and Nlrp3^-/-^ mice were stained with hematoxylin-eosin to visualize inflammatory cells (shown by an arrow). The tissue sections were scored in a blinded manner using the specific criteria listed in the material and methods. The inflammation scores for individual bladder (B) and kidney (D) samples were presented as a scatter plot with median as the central tendency. Statistical significance was determined by Mann-Whitney U test.

## Discussion

Various reports have established a crucial role for NLRP3 inflammasome in acute and chronic cystitis caused by uropathogenic *E. coli* (UPEC) (15-23). Whether NLRP3 inflammasome is important in MRSA-UTI, however has not been deciphered. In this report, we sought to bridge this knowledge gap by comparing MRSA UTI between C57BL6 WT and *Nlrp3*^-/-^ mice carrying a targeted mutation in *Nlrp3* gene (30). As a model organism, we used uropathogenic MRSA 1369 that has been previously used in UTI research (12, 13). We have also reported that *in vitro* exposure to human urine induces MRSA 1369 α-hemolysin (13), a known activator of NLRP3 in myelocytes (25). We observed consistent but statistically insignificant reduction in the CFUs recovered from the bladder and kidneys of *Nlrp3*^-/-^ mice compared to the WT at 24 and 72 h after the induction of ascending UTI. Moreover, the number of mice from *Nlrp3*^-/-^ cohort with detectable MRSA burden in kidneys was also lower compared to the WT. In contrast to these results, in the mouse model of CAUTI, the median MRSA CFUs recovered from the bladder and the kidneys, the overall spread of the data around median, and the number of mice with detectable CFU burden were alike between the WT and *Nlrp3*^-/-^ backgrounds at 24 and 72 hpi. Examining the MRSA infection in WT and *Nlrp3*^-/-^ mouse models of CAUTI is clinically relevant because the use of indwelling urinary catheter is a known risk factor for MRSA-UTI (7). The administration of NLRP3 inhibitor MCC950, 4 hours after the induction of ascending UTI without catheter, resulted in a 3-fold, statistically significant reduction in the bladder CFUs at 24 hpi and a corresponding reduction in the kidney CFUs that was statistically non-significant. Compared to the DMSO-treated controls, fewer MCC950-treated mice showed kidney CFUs. MCC950 is a small molecular inhibitor of selectively inhibits the formation of NLRP3 inflammasome complex by binding the NLRP3 NACHT domain and blocking ATP hydrolysis (35, 36). In addition, the immune profiling of mice also revealed that NLRP3 activity does not affect either the cytokine production or immune cell infiltration in the MRSA infected bladder and kidney tissues. The histopathological examination of bladder and kidney sections also showed that MRSA 1369 induced inflammation was not significantly altered in *Nlrp3*^-/-^ mice. Overall, these results argue that NLRP3 activity may be largely dispensable in MRSA-UTI. This is different from UPEC experiments where NLRP3-deficient (*Nlrp3*^-/-^ and *ASC*^-/-^) mice showed more severe acute cystitis marked by significantly higher UPEC burden, neutrophil influx, and inflammatory pathology in the infected mouse bladder (37).

The similar IL-1β levels in MRSA 1369-infected *Nlrp3*^-/-^ and WT mice suggested that IL-1β may be processed in an NLRP3 inflammasome independent manner similar to previous report that in neutrophils infected with UPEC strain CFT073, pro-IL1β to IL-1β processing was mediated by a cytoplasmic serine protease (20). It has been previously reported that MRSA 1369 infection exacerbates catheterization-induced localized inflammation at 24 hpi (12). The bladder and the kidneys from mice infected without catheter implant with MRSA strain SA116 also showed higher levels of pro-inflammatory cytokines, IL-1β, IL-6, and TNFα at 24 hpi (38). In contrast, we observed that MRSA 1369 mediates a modest increase in the levels of various pro-inflammatory cytokines and chemokines at 24 hpi. This difference may be attributed either to the presence of catheter implant (12), or to the potential differences between SA116 and MRSA 1369 (38).

Overall, our results argue a minor role, if any, for NLRP3 in shaping urinary immune defenses against acute MRSA UTI. Since we examined infection parameters in the WT and *Nlrp3*^-/-^ mice up to 72 h after the induction of ascending UTI, future experiments focused on determining whether NLRP3 activity plays a role in the chronicity and life-threatening exacerbations of MRSA UTI are warranted.

